# Modular Inter-brain Synchrony Network Associated with Social Difficulty in Autism Spectrum Disorder: a Graph Neural Network-Driven Hyperscanning Study

**DOI:** 10.1101/2025.06.18.660087

**Authors:** Yuhang Li, Yiting Zhu, Yingbo Geng, Danyong Feng, Shuo Guan, Dongyun Li, Lianni Mei, Xingyu Ding, Yupeng Ying, Jiaxuan Tang, Jiacheng Liang, Yueting Su, Yingchun Zhang, Qiong Xu, Rihui Li

## Abstract

Understanding social difficulties in Autism Spectrum Disorder (ASD) remains challenging due to its neurobiological heterogeneity and the limited ecological validity of conventional neuroimaging methods in capturing dynamic social interactions. Hyperscanning analysis based on functional near-infrared spectroscopy (fNIRS), which measures inter-brain synchrony (IBS) during dyadic interaction, offers a novel avenue to address these challenges. However, prior studies on ASD have reported inconsistent findings, primarily focusing on intra-regional synchronization while overlooking cross-regional network dynamics. To bridge this gap, we proposed an interpretable graph neural network (GNN) model to systematically identify ASD-specific IBS modular network between child-caregiver dyads during naturalistic cooperative puzzle-solving and free-talking tasks. We identified distinctive key IBS sub-networks for the cooperative puzzle-solving task and free-talking task, with the frontal eye field (FEF) of caregivers, the dorsal lateral prefrontal cortex (DLPFC) and the motor region of children highlighted. Furthermore, the key IBS sub-networks were found to be able to predict multiple domains of the core ASD symptoms. By integrating hyperscanning with GNN-driven analysis, this work uncovers task-dependent inter-brain neural mechanisms underlying social difficulties in ASD. These findings advance the field by proposing a data-driven framework to identify IBS biomarkers tied to clinical profiles, paving the way for personalized interventions that integrate computational neuroscience with clinical practice.

## Introduction

Autism Spectrum Disorder (ASD) is a prevalent and heterogeneous neurodevelopmental condition characterized by persistent deficits in social communication and interaction, alongside restricted and repetitive behaviors, interests, or activities (1,2). Globally, ASD affects approximately 1 in 100 individuals (3), with a reported prevalence of 1 in 142 in China (4). The rising prevalence of ASD, coupled with the lifelong need for adaptive support, imposes substantial socioeconomic burdens on families and healthcare systems (5). Despite its profound impact, accurately characterizing core autistic symptoms (e.g., social interaction abilities) remains a significant challenge due to the heterogeneity of behavioral phenotypes in ASD (6,7), hindering early diagnosis, precise clinical assessment, and reliable prognostic evaluation. Consequently, elucidating the neural mechanisms underlying social deficits in individuals with ASD is imperative to enhance diagnostic accuracy and guide the development of effective interventions.

Advances in neuroimaging techniques, including functional magnetic resonance imaging (fMRI), functional near-infrared spectroscopy (fNIRS), and electroencephalography (EEG), have identified atypical activation patterns in ASD individuals during social tasks, particularly within key regions of the “social brain”, such as medial temporal gyrus (MTG), temporoparietal junction (TPJ), and superior temporal sulcus (STS) (8–12). However, these findings predominantly derive from single-brain paradigms that isolate participants in artificial laboratory settings (e.g., viewing static social stimuli on screens). This methodological limitation restricts the ecological validity given that real-life communication requires individuals to actively engage in behavioral coordination based on real-time feedback from interaction partners, rather than passively responding to fixed stimuli. By isolating participants and using static stimuli, conventional paradigms could overlook the mechanisms (e.g., predictive coding, mutual adaptation) that underpin the social difficulty that individuals with ASD face during daily life (13).

Hyperscanning, a technique that enables the simultaneous recording of brain activity from multiple individuals, has emerged as a tool for studying social cognition in naturalistic contexts with real-life social exchange (14–17). By quantifying inter-brain synchrony (IBS), the neural coupling between interacting partners, hyperscanning provides a method to delineate the interpersonal neural mechanisms during social interaction (17–19). Studies have demonstrated that IBS reflects both the quality of interaction (e.g., mutual engagement, reciprocity) and the relational dynamics between individuals (20,21). These findings underscore IBS as a neural marker of successful social coordination, highlighting its potential to elucidate social impairments in ASD (22).

However, prior hyperscanning investigations on the social interaction of ASD dyads revealed inconsistent evidence. While some studies reported diminished IBS between children with ASD and experimenters compared to typically developing (TD) dyads (23), others found no group differences despite observable behavioral synchrony (24,25). One of the possible reasons for the lack of group differences observed in some studies could be the absence of face-to-face interaction between the dyads in their experiments (24,25). Moreover, previous studies investigating IBS in ASD during social interactions have predominantly focused on intra-regional synchronization patterns (e.g., the IBS between the right TPJs of parents and children), overlooking the critical role of cross-regional neural synchrony. Recent methodological advancements address this limitation by constructing IBS networks that quantify synchronization across all possible channel pairs between interacting brains (19,26). This approach offers a more comprehensive characterization of dyadic social dynamics, capturing both localized and distributed neural coupling mechanisms. Yet, a systemic, network-based framework remains to be explored for assessing how and whether IBS networks between children with ASD and their caregivers differ from TD dyads during social interactions, thereby unraveling the neurobiological underpinnings of social impairments in ASD.

Graph Neural Network (GNN) has emerged as a transformative tool for analyzing complex brain networks, offering unique capabilities to model non-Euclidean connectivity patterns inherent to neuroimaging data (27–29). Unlike traditional methods constrained by grid-based architectures (e.g., CNNs), GNNs explicitly encode relationships between nodes (brain regions) and edges (functional/structural connections), enabling the extraction of topological features critical for understanding neurodevelopmental disorders (30). In single-brain analyses, GNNs have demonstrated remarkable success in identifying disorder-specific biomarkers, such as dysregulated connectomes in schizophrenia (31) and altered sub-networks in ASD structural networks (32). Crucially, recent advances in interpretable GNNs further enhance their utility in clinical research (33–35). Thereafter, interpretable GNNs hold particular promises for unraveling the atypical IBS network pattern of the ASD dyads.

This study aimed to investigate distinctive IBS network patterns between children with ASD and their caregivers compared to TD dyads by leveraging an interpretable GNN. Children and their caregivers participated in two naturalistic face-to-face social interaction tasks (puzzle-solving and free-talking), during which their hemodynamic activities were recorded using a fNIRS system. We trained an interpretable GNN model on the derived IBS networks to classify ASD and TD groups, identifying key discriminative sub-networks through GNN-generated explanation masks. Furthermore, we evaluated associations between these sub-networks and clinical symptom severity in ASD. By integrating GNNs into IBS analysis, this approach offers novel insights into the neural mechanisms underlying social impairments in children with ASD.

## 2. Method

### 2.1 Participants

We recruited 44 children with ASD (age 7.10 ± 2.02 years, 6 females) and their caregivers from the Children’s Hospital of Fudan University, Shanghai, China. The diagnosis of ASD was conducted by specialist pediatricians from the hospital using rigorous diagnostic criteria specified in the Diagnostic and Statistical Manual of Mental Disorders, 5th edition (DSM-5) (1), following more than 30 minutes of direct clinical observation, supplemented by the Autism Diagnostic Observation Schedule (ADOS). Exclusion criteria included (1) ASD with a known genetic condition (e.g., *MECP2*, *TSC1* or *TSC2*-related neurodevelopmental disorder); (2) ASD with a known neurological medical condition in the acute phase (e.g., epilepsy, head traumas, encephalitis, or meningitis). Besides, we recruited 48 age- and sex-matched TD children (age 7.90 ± 2.28 years, 13 females) and one of their caregivers.

### 2.2 Behavioral assessment

For each child with ASD, their clinical symptoms were assessed using the Autism Diagnostic Observation Schedule, Second Edition (ADOS-2) (36), which evaluates social communication skills and restricted and repetitive behaviors. Cognitive abilities were measured using the Chinese version of the Wechsler Intelligence Scale for Children-Fourth Edition (WISC-IV) (37). The caregivers of children in the ASD group completed a series of behavioral assessments on the same day as neuroimaging data collection, including Vineland Adaptive Behavior Scales, Second Edition (Vineland-2) (38,39) , and Behavior Rating Inventory of Executive Function, Second Edition (BRIEF-2) (40). The Vineland-2 provides a nuanced understanding of an individual’s functional capabilities and needs. It assesses an individual’s ability to function independently and adapt to their environment by various sub-domains, including communication, daily living skills, socialization, motor skills, and overall level of adaptive functioning. BRIEF-2 measures the children’s development of executive functions in 4 sub-domains: Behavior Regulation Index (BRI), Emotion Regulation Index (ERI), Cognitive Regulation Index (CRI), and Global Executive Composite (GEC).

### 2.3. Experiment procedure

After the participants’ visit, children and their caregivers were instructed to sit on opposite sides of a table face-to-face. They were then introduced to the experiment procedure, as shown in Figure 1A, while wearing the brain recording devices. The experiment consisted of 3 parts: puzzle-solving, free-talking, and resting (Figure 1A). All participants completed the tasks in this order. The time intervals between tasks were at least 30 seconds. During the interval, the experimenters introduced the next task again and emphasized the requirements.

**Figure 1.**
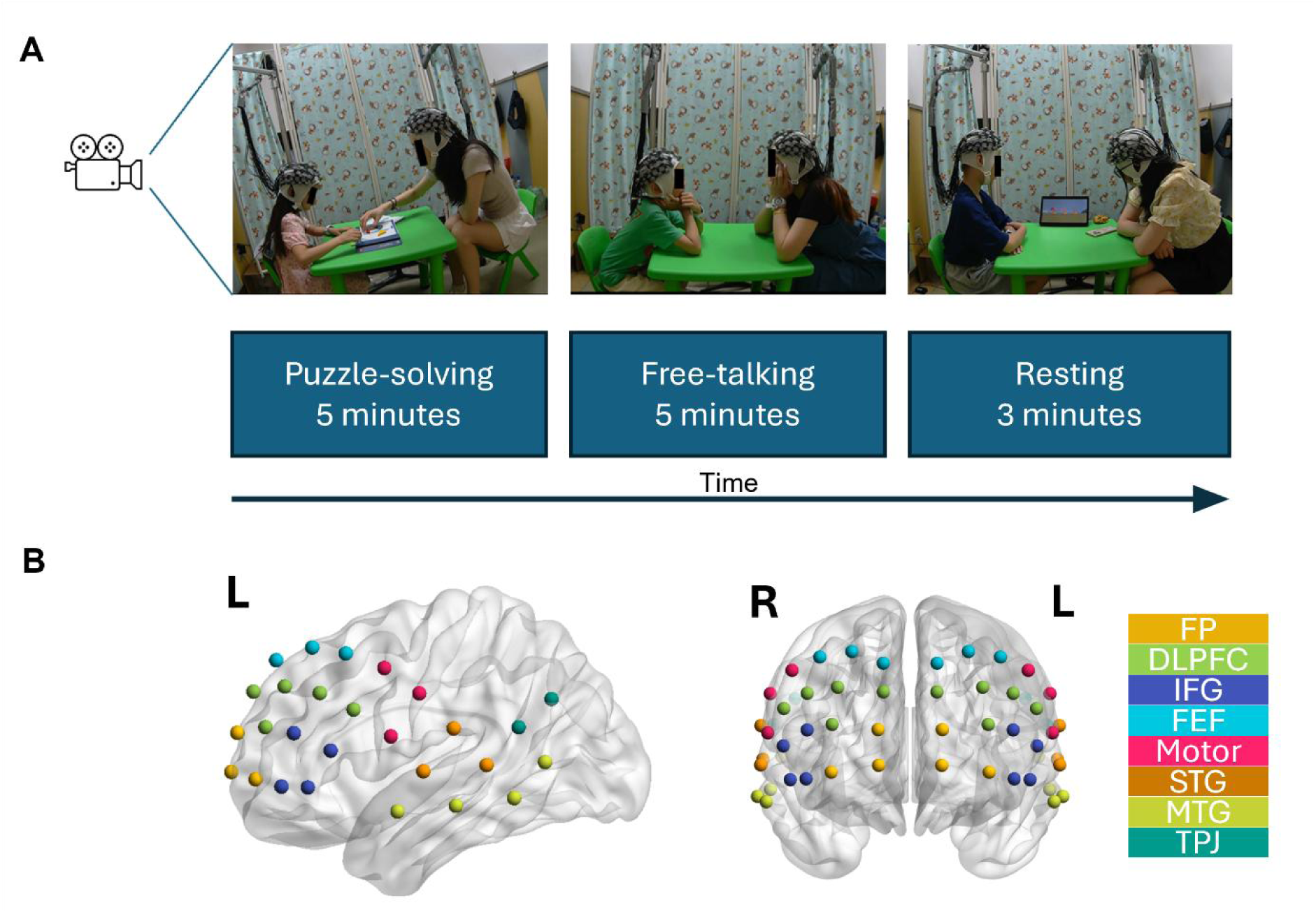
The experimental settings. (A) The experimental paradigm. The participants were instructed to complete three stages of experiments, including puzzle-solving, free-talking and resting, while their interactions were recorded by camera. (B) The arrangement of fNIRS channels. The channels projected on the same ROI are in the same color as labeled in the right panel.

#### Puzzle-solving task

Children and their caregivers were instructed to cooperatively complete as many jigsaw puzzles as possible within a 5-minute time frame. Each puzzle shows a picture of a target shape composed of 7 pieces. The participants’ goal was to put the 7 pieces together correctly. To better engage the children in the task, the experimenters provided participants with puzzles of 3 difficulty levels and chose the puzzles for the formal experiment according to the children’s interest.

#### Free-talking task

Children and their caregivers were instructed to talk with each other about topics for 5 minutes. We provide two topics for their choices: 1) The last time they went out for fun together; 2) The plan for their next vacation.

#### Post-task Resting

Children and the caregiver were required to sit still with no conversation for 3 minutes. To facilitate compliance and help the children remain calm, a short cartoon episode was played during this period.

### 2.4. Data Acquisition of fNIRS

The hemodynamic activities of both children and their caregivers were recorded simultaneously with a continuous-wave fNIRS system (Danyang Huichuang, China) with 24 sources and 24 detectors for each of them at a sampling rate of 11 Hz. The same channel arrangements covered the bilateral frontal and temporal regions of both children and adult caregivers (Figure 1B). A digital video camera was placed beside the participants. Both the video and the audio were recorded.

### 2.5 Data analysis

**2.5.1 Inter-brain Synchrony (IBS) network construction**

We calculated the wavelet transform coherence (WTC) for each pair of possible channels with the *wcoherence* function in MATLAB Signal Processing Toolbox (41). The WTC values were further Fisher transformed to z scores. We then averaged the WTC matrix (time × frequency) across time for each task. To identify the frequency of interest, we compared the IBS of puzzle-solving and free-talking tasks against the resting task by pair-wise t-tests. The frequency bins that showed significantly higher IBS during social interaction above resting were selected as the frequency of interest. The significances were corrected with the FDR method across all channel pairs. After averaging across all the selected frequency bins (puzzle-solving: 0.184∼0.292 Hz; free-talking 0.184∼0.260 Hz), the WTC values of all possible channel pairs formed the IBS network for each dyad (Figure 2B).

**Figure 2.**
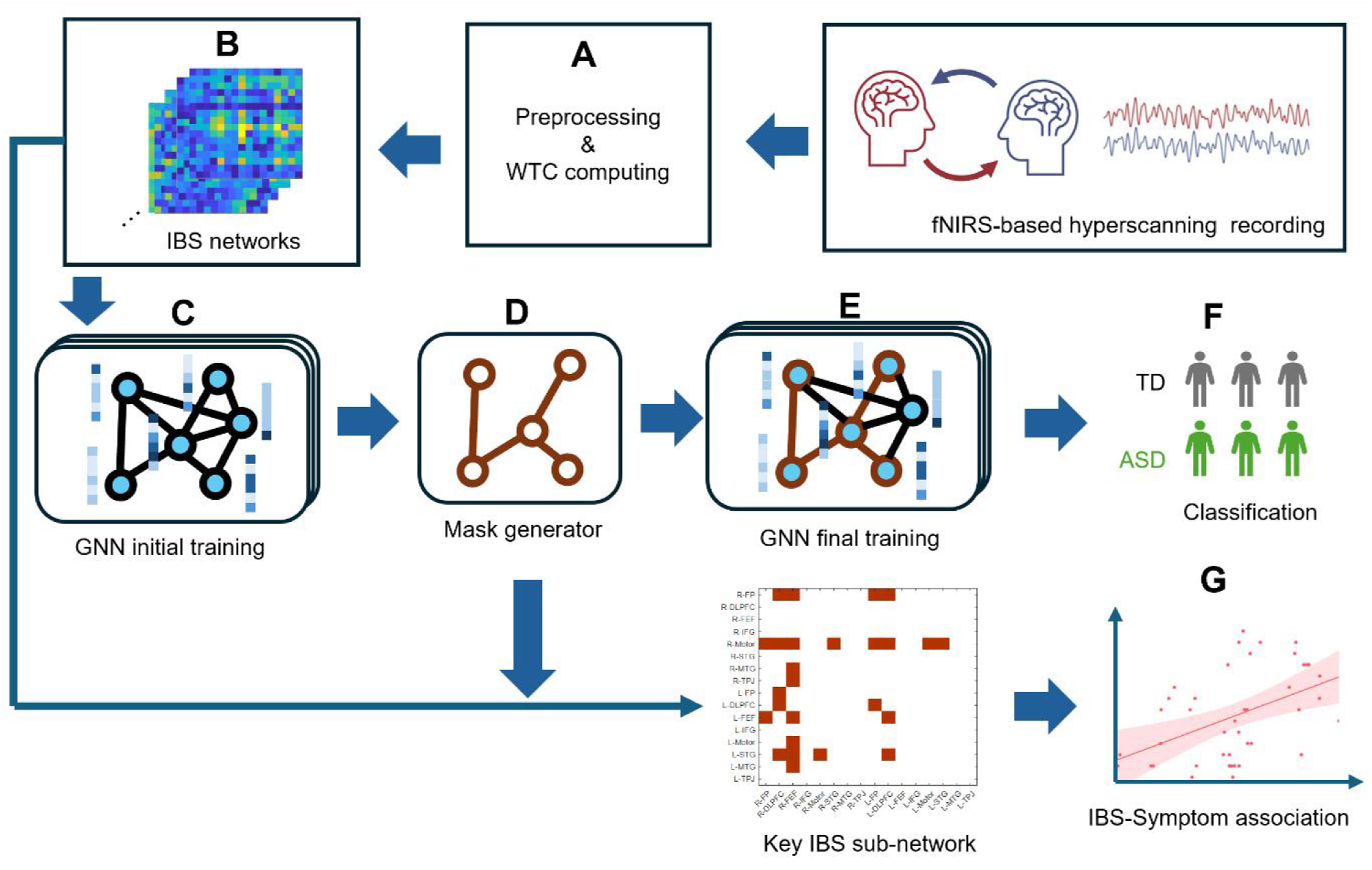
The data analysis pipeline. The IBS networks were computed from the preprocessed fNIRS signals recorded simultaneously. Then, a GNN model was implemented to classify the IBS networks between the ASD and TD groups while generating a weighted mask. The IBS networks were filtered with the mask to get key IBS networks, which were input to predict the clinical symptoms of children with ASD.

#### 2.5.2 GNN-driven Inter-brain Synchrony Network

As shown in Figure 2, an interpretable GNN model was implemented to classify the ASD and TD groups by their IBS networks, from which the key IBS network for achieving the best classification was extracted (33). Briefly, the GNN model was trained on node features and edge weights of all the samples and predicted the category of each sample. The IBS network between each pair of participants can be regarded as an asymmetric graph. Each channel is a node of the graph, and the IBS between channels represents the edge weights. For each node, its node features were a vector concatenating the edge weights between itself and all other nodes. After the initial training with multiple GNN layers, the model trained a matrix of weights as the mask of the IBS network input (Figure 2C-D). Finally, the GNN model was trained with the masked IBS network input and generated the model prediction (Figure 2E-F). Hyperparameters were selected with the open-source AutoML toolkit NNI (42). The model accuracy results were averaged across five-fold validation.

The same training procedure was performed for the puzzle-solving task and the free-talking task separately. The best-fitted models were finally trained on all samples and generated masks of IBS networks for both tasks. Then the channel masks were merged into ROI masks by averaging the weights of channels within each ROI. The ROI pairs with top 10% mask weights were selected as key IBS sub-networks in the classification between the ASD and TD groups.

#### 2.5.3 Encoding of social interaction behaviors during puzzle-solving

The social interaction behaviors of both children and caregivers during the puzzle-solving task were encoded into 18 categories (See supplement materials Table S2) according to the dyadic parent-child Interaction Coding System (DPICS) (43). Two research assistants conducted the behavior encoding independently and recorded the times each behavior category occurred during the task. The final frequency of each behavior category during the 5-minute task was averaged across two research assistants.

#### 2.5.4 Conversation analysis

To evaluate the speech task performance before and after learning, the speech recording was first transcribed using the Computerized Language Analysis (CLAN) (44). In alignment with the IELTS speaking rubrics, the speech performance was quantitatively evaluated from fluency and coherence, and lexical and grammatical complexity (45,46). 14 features of language characteristics were generated from the conversation of each dyad (See supplement materials Table S3).

### 2.6 Statistical analysise

Within the key IBS sub-network, nodal strengths of each ROI were computed by summing the IBS values between each ROI and all other ROIs. We conducted independent sample t-tests to compare the node strengths of the ASD group against the TD group. False detection ratio (FDR) correction was implemented with the Benjamini-Hochberg method. To investigate the association between the key IBS sub-network and core ASD symptoms, we employed Lasso regression with leave-one-out cross-validation (Figure 2F). The independent variables comprised IBS values from key IBS sub-networks, while dependent variables were the scale scores of ASD symptoms, including all the sub-domains of WICS-IV, ADOS-2, Vineland-2, and BRIEF-2. The regularization strength λ, which determines weight sparsity through coefficient shrinkage, was optimized via logarithmic grid search across 100 candidate values with range from 0.0001 to 1. Model performance was evaluated by computing Pearson’s correlation coefficient between predicted and observed symptom scores.

## 3 Results

### 3.1 Demographic characteristics of participants

The demographic characteristics of all participants are summarized and shown in Table 1. There was no significant difference in age (*p* = 0.64) and sex (*p* = 0.11) between the two groups.

**Table 1.**
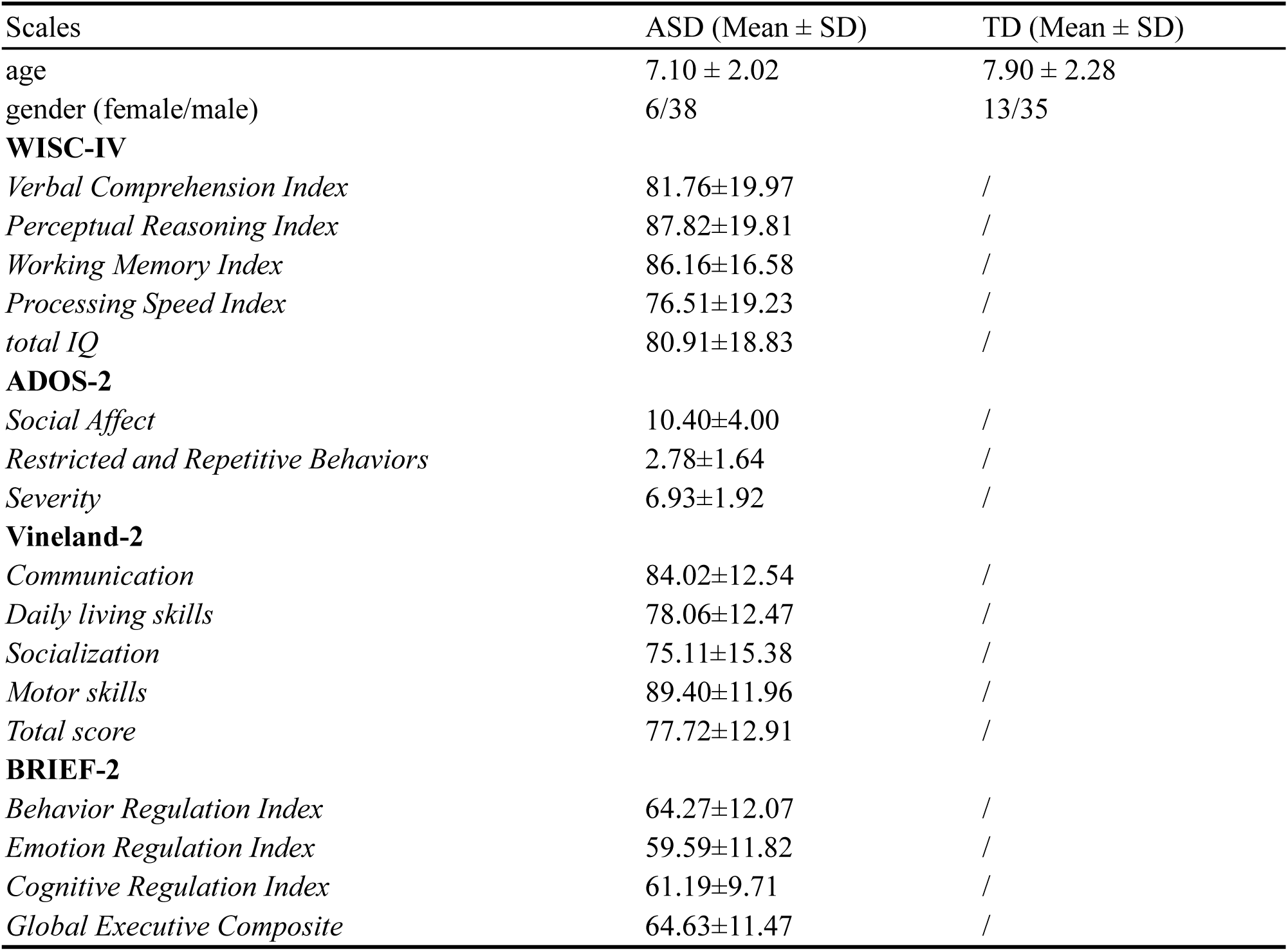
Clinical characteristics of children. TD, typical developed; ASD, autism spectrum disorder

### 3.2 Atypical behavioral patterns and language characteristics of ASD

Compared with those of TD children, the frequency of “No answer” and “No comply” was higher for children with ASD during the cooperative puzzle-solving task (t(_91_) = −5.02, *p* <.001; (t(_91_) = −5.24, *p* < .001;). The TD children asked questions (t(_91_) = 8.95, *p* < .001), proposed talks (t(_91_) = 3.39, *p* = .001), complied with commands from the caregiver (t(_91_) = 3.26, *p* = .002), and initiated negative talks (t(_91_) = 4.14, *p* < .001) with higher frequency during the tasks (Figure 3A).

**Figure 3.**
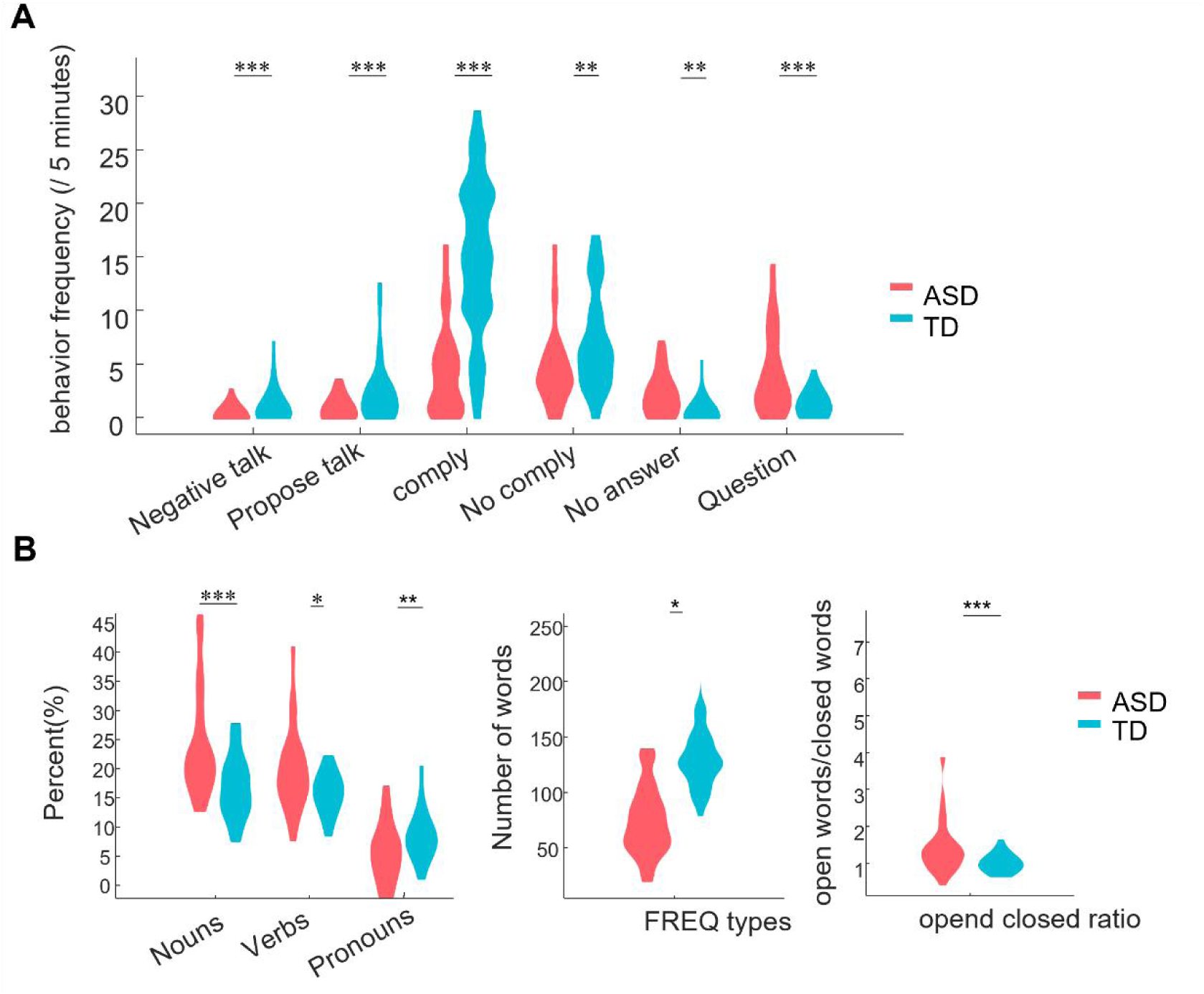
The group comparison of the coded behavioral patterns and the children’s language characteristics. (A) The comparison of coded dyadic behaviors during the puzzle-solving task. (B) The children’s language characteristics during the free-talking task. All significant differences were FDR corrected. *: p <.05; **: p <.01; ***: p < .001

The children with ASD showed distinctive language characteristics during the free-talking task. Children with ASD used more verbs (t(_91_) = 3.88, *p* = .001) and nouns (t_(91)_ = 4.78, *p* < .001) but less pronouns (t_(91)_ = - 2.06, *p* = .031). Moreover, the FREQ types of ASD group were lower than TD group (t_(91)_ = −3.05, *p* = .015). Finally, children in the ASD group had a significantly higher open-close ratio in their word choices during talking (t_(91)_ = 4.12, *p* < .001) (Figure 3B).

### 3.3 Inter-brain synchrony network identified by Graph Neural Network

The trained GNN model achieved 75.2% accuracy for the puzzle-solving task and 72.3% accuracy for the free-talking task. The model generated a weighted mask of the IBS networks for each task separately. We selected the ROI pairs with top 10% weights as the key IBS sub-networks for distinguishing the ASD and TD groups.

For the puzzle-solving task, the right frontal eye field (FEF) and right DLPFC of the caregivers and the right motor region of the children revealed more connections with other brain regions, indicating that these regions played a critical role in distinguishing the ASD and TD groups (Figure 4). The other brain regions in the key sub-network that contribute to the classification included the children’s left middle temporal gyrus (MTG) and the right frontal polar (FP), and the caregivers’ left DLPFC.

**Figure 4.**
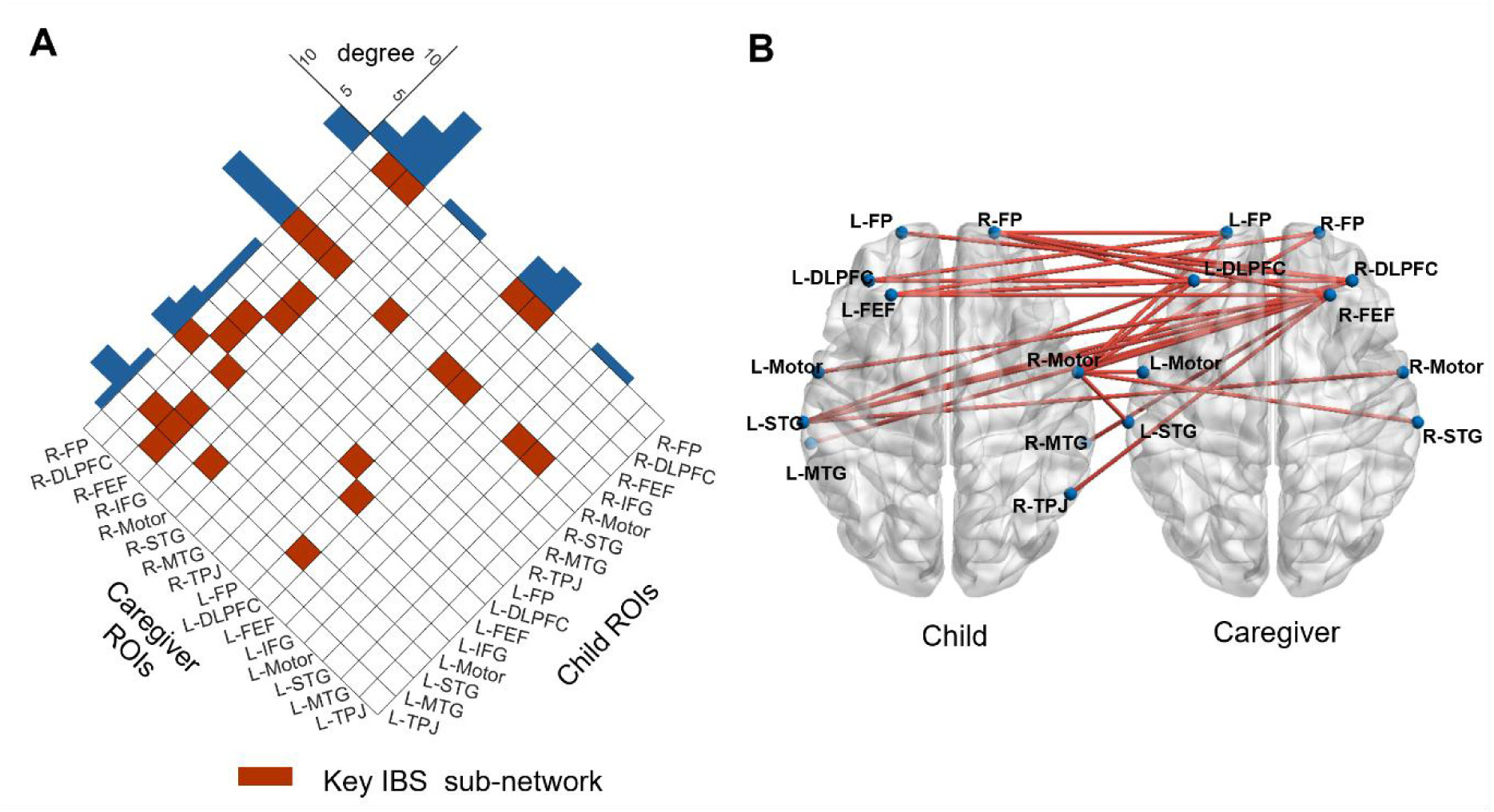
The ASD-specific key sub-network during the puzzle-solving task was identified by the GNN model. (A)The sub-network for the puzzle-solving task. The ROI pairs marked with deep red are selected key IBS sub-networks (top 10%). The degrees of ROIs are shown by the blue bars on the top. (B) The key IBS sub-network is shown on the inter-brain.

For the free-talking task, the left DLPFC of the children and the left FEF of the caregivers played a critical role in the key IBS sub-network (Figure 5). In addition, the right TPJs of both the child and the caregiver are also important brain regions involved in the key IBS sub-network.

**Figure 5.**
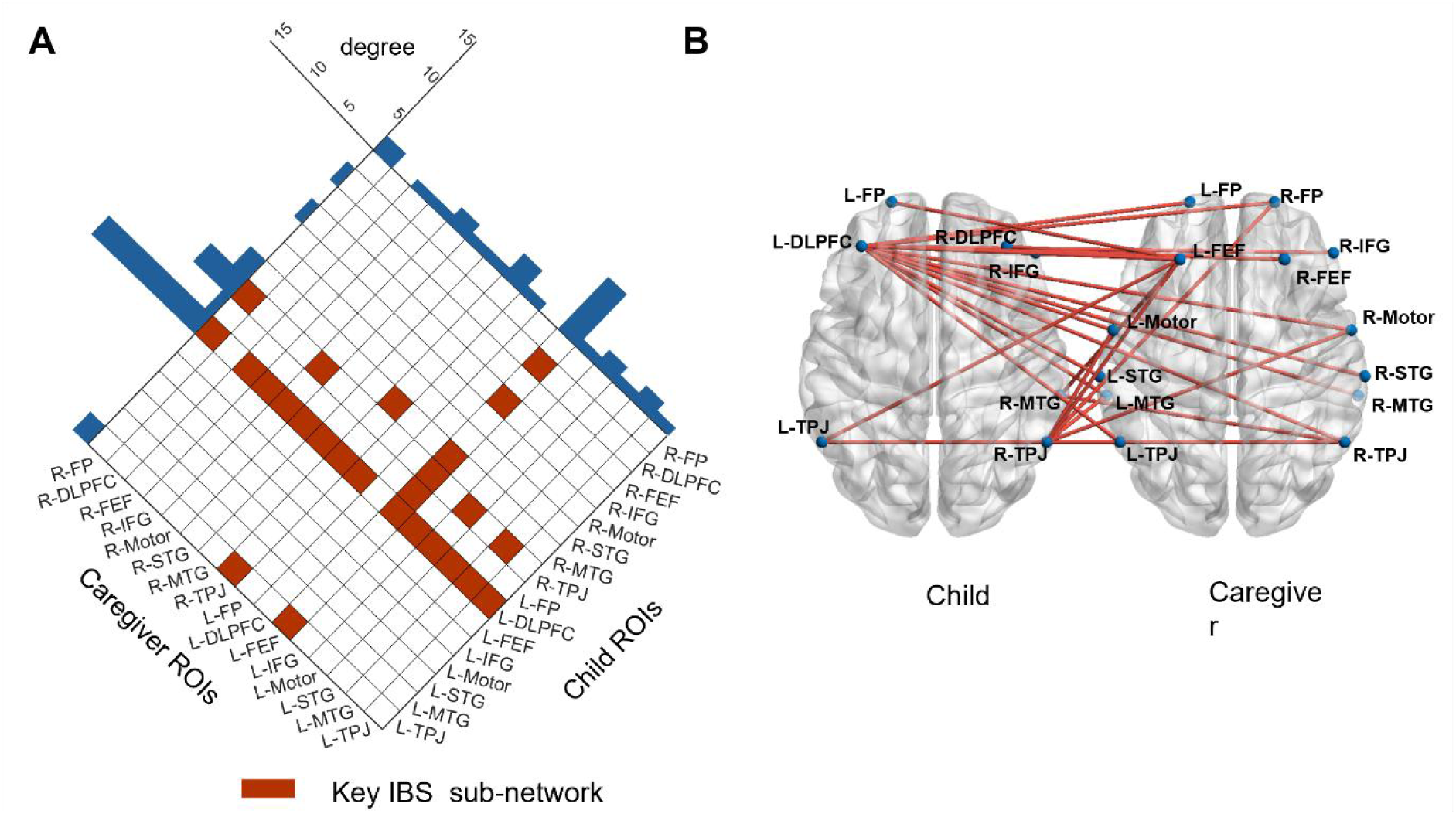
The ASD-specific key sub-network during the free-talking task was identified by the GNN model. (A)The weighted mask for the free-talking task. The ROI pairs marked with deep red are selected key IBS sub-networks (top 10%). The degrees of ROIs are shown by the blue bars on the top. (B) The key IBS sub-network is shown on the brains.

### 3.4 Prediction of the ASD related symptoms with key IBS sub-networks

We found that scores of 5 sub-domains could be predicted by the key IBS sub-network extracted from the puzzle-solving task using Lasso regression. These scales included the restricted and repetitive behavior (RRB) measured by ADOS-2 (*r* =.70, *p* < .001), the Communication (*r* = .38, *p* = .040), Socialization (*r* = .39, *p* = .035), and Total Score (*r* = .53, *p* = .002) of Vineland-2, and BRI (*r* =.43, *p* = .027) measured with BRIEF-2. The sparsified model weights reflected the critical nodes within the sub-network that contribute substantially to the model’s predictive capabilities. A clear divergence was observed among the salient nodes associated with the five subscales (Figure 6). For the RRB assessed with ADOS-2, most of the ROI pairs within the key IBS sub-network significantly contribute to the model prediction (Figure 6A). For the Socialization, Communication, and Total Score domains assessed using Vineland-2, convergent salient node distributions were found in the key IBS sub-network (Figure 6B-D). These nodes were predominantly localized to the right DLPFC and right FEF of the caregivers, demonstrating statistically significant positive weight coefficients. For the Behavioral Regulation assessed with BRIEF-2, only four significant ROI pairs were identified, including three negative weights associated with the right FEF of the caregivers (Figure 6E).

**Figure 6.**
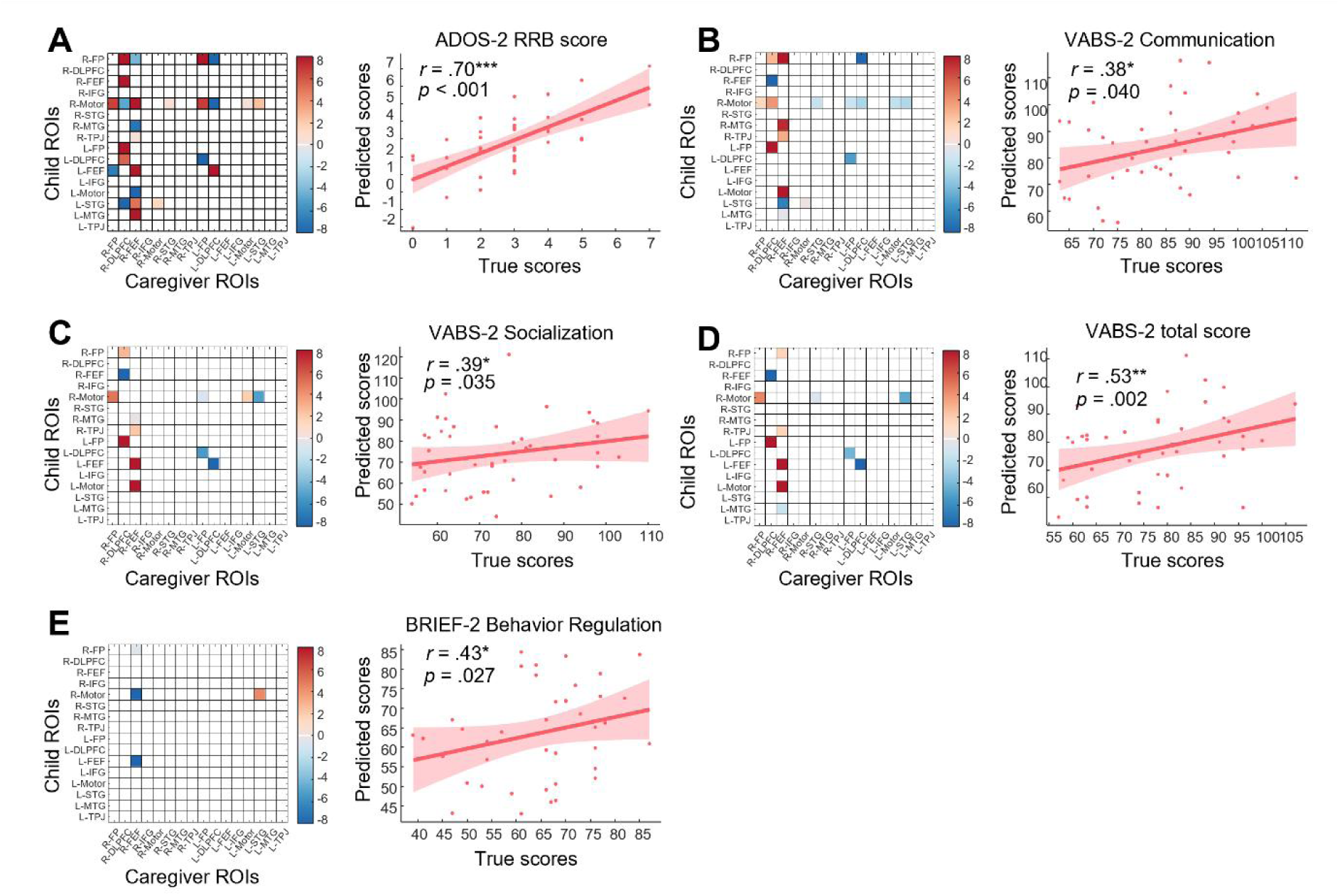
Prediction of the ASD related symptoms with the key IBS sub-network of puzzle-solving task. The model weights and the prediction performance of sub-domains that were significantly predicted with key IBS sub-network of puzzle-solving task, including (A) Restricted and repetitive behavior measured by ADOS-2, (B) Communication measured by Vineland-2, (C) Socialization measured by Vineland-2, (D) Total score of the Vineland-2, and (E) Behavior regulation measured by BRIEF-2.

For the free-talking task, we found that the RRB measured by ADOS-2 (*r* =.51, *p* = .011) (Figure 7A) and the Emotional Regulation (*r* =.47, *p* = .017) (Figure 7B) assessed by BRIEF-2 could be predicted by the key IBS sub-network. While the relatively sparse salient weights were found in predicting the RRB, most of the ROI pairs within the key IBS sub-network contributed to predicting the ERI (Figure 7).

**Figure 7.**
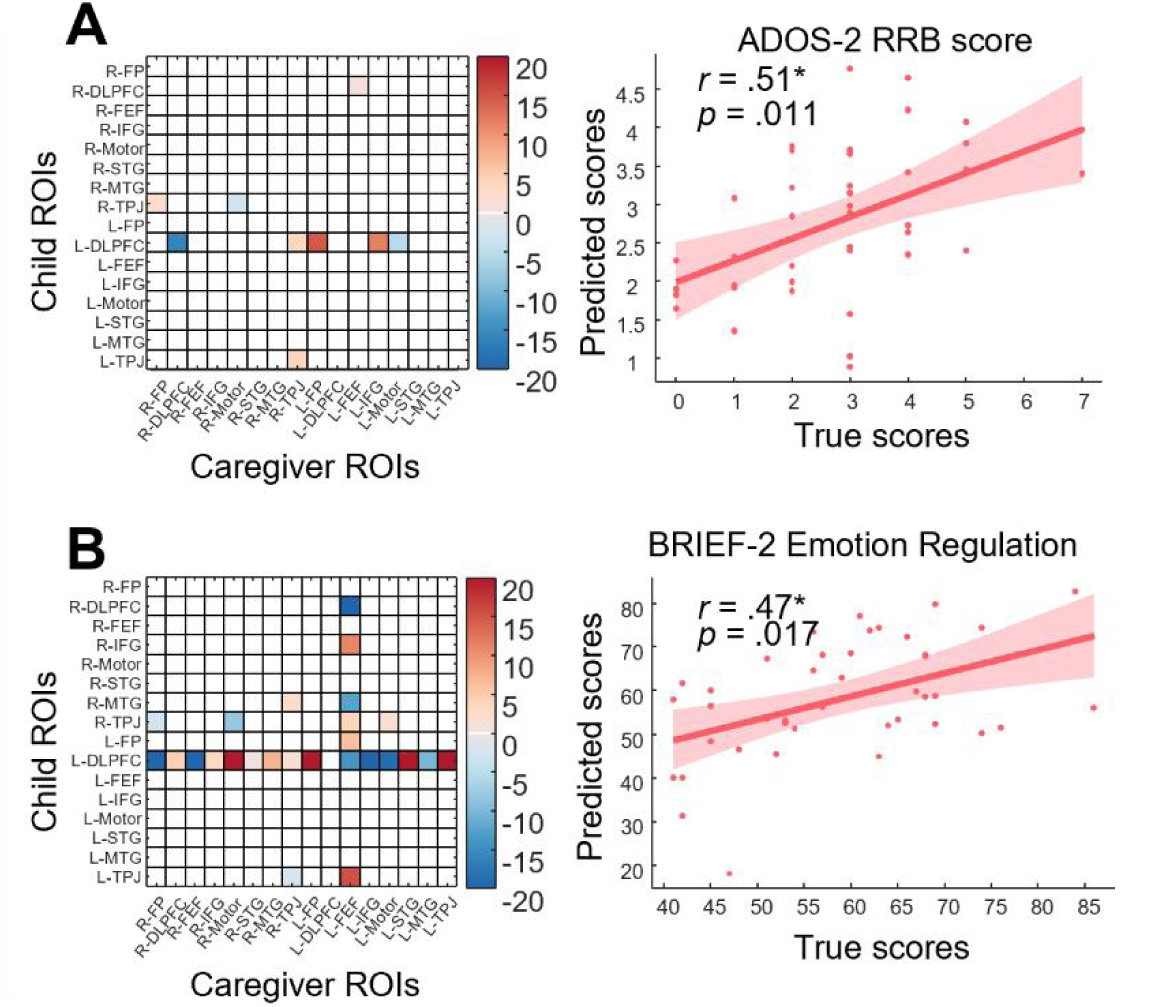
Prediction of the ASD related symptoms with key IBS sub-network during free-talking task. The model weights and the prediction performance of sub-domains that were significantly predicted with the key IBS sub-network of the free-talking task, including (A) The restricted and repetitive behavior measured by ADOS-2, and (B) Emotion regulation measured by BRIEF-2.

## 4. Discussion

Understanding of the neural mechanisms underlying social dysfunction in autism spectrum disorder (ASD) has long been hampered by its clinical and neurobiological heterogeneity, as well as the low ecological validity of traditional neuroimaging paradigms in capturing real-world social interactions. To address these gaps, this study integrated naturalistic tasks with interpretable GNNs to identify ASD-specific IBS network patterns between children and caregivers. We found that, during the puzzle-solving task, the distinctive IBS pattern of the ASD group was captured by a modular IBS sub-network involving the right FEF and the right DLPFC of caregivers and the motor regions of children. In contrast, during the free-talking task, IBS between the left DLPFC of children and the caregivers’ left FEF played a critical role in the classification of the ASD and TD groups. Moreover, the autistic symptoms assessed by ADOS-2, Vineland-2, and BRIEF-2 were found to be predicted by the IBS of the key IBS sub-network. The integration of GNN explanation masks establishes a methodological framework to objectively identify neural biomarkers tied to behavioral phenotypes, bridging computational neuroscience with clinical phenotypes. These findings provide a foundation for mechanistically grounded interventions targeting dyadic neural alignment in ASD.

### 4.1 Implementation of the GNN model in the IBS analysis

Our study introduces methodological advancements in hyperscanning research through GNN-based analysis of IBS networks. Unlike conventional approaches limited to isolated region-to-region synchrony analysis in ASD research (24,47,48), the proposed GNN framework captures task-dependent IBS patterns by modeling distributed, non-Euclidean network topology inherent in dynamic social interactions (49). The model’s interpretability, enhanced through explanation masks (33), enables the identification of critical sub-networks associated with behavioral phenotypes, which facilitates the discovery of neural biomarkers for social impairment disorders.

Through GNN-generated interpretable masks, we identified social interaction-specific IBS sub-networks in ASD dyads. During the puzzle-solving task, the key IBS sub-network involved the neural coupling between the caregiver’s right DLPFC and FEF and the child’s right motor region. In the free-talking task, critical neural coupling emerged in the child’s left DLPFC and the caregiver’s left FEF. Specifically, the involvement of the caregiver’s FEF across both tasks revealed distinct engagement patterns compared to TD dyads, potentially reflecting impaired joint attention mechanisms in ASD interactions (50–53). This aligns with evidence suggesting that caregivers of ASD children may experience challenges tracking their child’s attentional focus (54), manifesting as atypical IBS between caregiver FEF and children’s cortical regions. Children with ASD exhibited task-dependent neural signatures, such as the involvement of right motor during puzzle-solving versus left DLPFC engagement in free-talking. This dissociation likely reflects domain-specific social interaction impairments, in which motor regions support behavioral coordination in structured tasks while DLPFC mediates verbal exchange in social communication. These task-contingent patterns may originate from ASD-related restricted and repetitive behaviors disrupting social reciprocity, as evidenced by successful prediction of repetitive behaviors using IBS sub-networks (Figures 6A, 7A). Our network-level GNN analysis provides novel insights into ASD-associated IBS characteristics beyond traditional region-pair approaches. First, we reveal asymmetric neural disruptions in caregiver-child dyads, reflecting differential impacts of ASD symptoms on interaction partners. These findings advance understanding of social dysfunction mechanisms in ASD and could inform home-based intervention strategies. Second, we identified previously overlooked network features, particularly caregiver FEF involvement, a novel finding in ASD hyperscanning literature, suggesting disrupted social attention dynamics. Collectively, our GNN framework establishes a transformative paradigm for hyperscanning research, offering scalable, interpretable tools to decode IBS dynamics and accelerate biomarker discovery for neurodevelopmental disorders.

### 4.2 Prediction of the ASD symptoms: neural biomarker towards personalized treatment

Our study demonstrates that ASD symptoms measured by ADOS-2, Vineland-2, and BRIEF-2 can be predicted using IBS sub-networks derived from the interpretable GNN model. Critically, the RRB of ADOS-2 was predictable from IBS sub-networks extracted from both puzzle-solving and free-talking tasks. This extends beyond prior research, which primarily linked IBS patterns to social difficulties in ASD (48,55,56), by establishing a novel association between atypical IBS patterns in ASD dyads and children’s RRBs. Though successful predictions were obtained on both tasks, the underlying neural mechanisms differed. For the puzzle-solving task, the children’s right motor region and caregivers’ right FEF were highlighted in the prediction. This suggests that the children’s RRBs may trigger abnormal activation in motor regions (57), which subsequently impacts neural synchrony with the caregivers. Furthermore, during this task, the children’s RRBs likely made it more difficult for the caregivers to maintain typical social attention, causing the altered neural activity of the caregiver’s FEF region (58,59). Regarding the free-talking task, RRBs were predicted by a sparser IBS sub-network, with the children’s left DLPFC highlighted as a key region. Given the left DLPFC’s strong association with language functions (60,61), dysregulation in this region during the free-talking task likely contributes to the children’s restricted and repetitive speech patterns, which could disrupt the caregiver-child IBS network. Collectively, these findings demonstrate that GNN-based IBS network analysis not only advances mechanistic understanding of neural synchrony in ASD but also establishes a critical foundation for developing hyperscanning-driven diagnostic tools.

### 4.3 Behavioral and Linguistic Differences

Consistent with prior work, children with ASD exhibited significantly higher frequencies of “No answer” and “No-comply” during puzzle-solving tasks compared to TD children, aligning with theories of reduced social motivation (62) and observational studies of ASD dyads (2). A novel finding was the reduced initiation of negative talks by ASD children during free-talking, suggesting a core deficit in social spontaneity that warrants further exploration through neural synchrony analyses. The distinctive language patterns observed during the free-talking task further underscore the atypical social communication profile in ASD. Children with ASD used significantly more verbs and nouns but fewer pronouns, alongside reduced lexical diversity (i.e., lower FREQ types and a higher open-close word ratio). These findings align with prior reports of ASD-related language rigidity, including a preference for concrete, context-bound vocabulary over abstract or socially adaptive terms (63). The overuse of verbs and nouns may reflect compensatory strategies to anchor communication in tangible referents, circumventing challenges in perspective-taking and pronoun resolution inherent to social dialogue (64). Conversely, the scarcity of pronouns, a hallmark of pragmatic language impairment in ASD, may stem from difficulties in tracking conversational roles or inferring shared mental states (65).

### 4.4. Limitations and Future Directions

This study has several limitations that warrant consideration. First, while the sample size (*N* = 95 dyads) in the present study is relatively sufficient for a typical ASD study, it may limit the performance of our GNN model. Future studies should prioritize multicenter collaborations to recruit larger cohorts, enabling more robust GNN training and validation across diverse ASD subtypes. Second, the spatial resolution of fNIRS constrains our ability to resolve deeper subcortical structures implicated in social-emotional processing. Emerging techniques such as diffuse optical tomography (DOT), which enhances spatial specificity through dense optode arrays, could mitigate this limitation while retaining fNIRS’s advantages for naturalistic recording (66,67). Third, differences in parenting styles between ASD and TD caregivers may confound group-level IBS differences. While this introduces complexity, it also highlights the need to explore caregiver-child dynamics as a therapeutic target. Future work should incorporate standardized assessments of caregiver behavior to disentangle ASD-specific neural signatures from interactional adaptations.

## Acknowledgements

This work is supported by the National Natural Science Foundation of China (82301743), the Science and Technology Development Fund of the Macao SAR (0010/2023/ITP1, 0016/2024/RIB1), Guangdong Provincial Natural Science Foundation (2025A1515010539), the University of Macau (SRG2023-00015-ICI, MYRG-GRG2024-00296-ICI, and MYRG-CRG2024-00022-ICI), the Natural Science Foundation of Anhui Province (No. 2308085MH255), the China Medical Board Open Competition Program (Grant No. CMB-OC-22-471), and the academic leaders development program (EKXDPY202306) and Med+X cross-disciplinary team project of the Children’s Hospital of Fudan University.

## Disclosure

The authors declare no competing interests.

## Notes

### Competing Interest Statement

The authors have declared no competing interest.

