## Supplement material for "Modular Inter-brain Synchrony Network Associated with Social Difficulty in Autism Spectrum Disorder: a Graph Neural Network-Driven Hyperscanning Study"

1. **Preprocessing of fNIRS data**

The data analysis was conducted utilizing the NIRS Analyzer toolbox (1) within MATLAB alongside a customized MATLAB-based script. Initially, the raw fNIRS data was converted to optical density. Motion artifacts were then eliminated using a Temporal Derivative Distribution Repair (TDDR) method (2). The concentration changes of HbO and HbR were computed according to the Modified Beer-Lambert Law. The bad channels with no salient heartbeat signal were manually identified with time-frequency analysis and excluded from the following analysis. Principal component analysis (PCA) was used to remove global physiological noises, such as the skin blood flow (Zhang et al., 2005). The threshold for removing variance was set at the 90% conservative level. Finally, the signals were down-sampled to 4 Hz. Post-analysis in this study exclusively focused on HbO, given its recognized robustness and sensitivity compared to HbR regarding brain functional changes (R. Li et al., 2020; Plichta et al., 2006.). To delineate the regions of interest (ROIs) encompassed by the fNIRS channels, we projected all fNIRS channels onto the cortical surface, utilizing their MNI coordinates derived from the scalp surface employing an automatic anatomical labeling method. 16 ROIs covering bilateral frontal and temporal regions were included in the analysis. The MNI coordinates and brain regions of each ROI are listed in Tabel S1.

| **Table S1.** The covered ROIs. | | | | |
| --- | --- | --- | --- | --- |
| **MNI coordinates** | | | **Brain Area** | **Hemisphere** |
| **X** | **Y** | **Z** |  |  |
| 21.47 | 64.43 | 7.44 | Frontal polar | Right |
| 36.69 | 42.16 | 32.13 | Dorsal lateral prefrontal cortex | Right |
| 26.40 | 35.54 | 51.92 | Frontal eye field | Right |
| 51.17 | 38.81 | 6.15 | Inferior frontal gyrus | Right |
| 60.20 | -0.25 | 33.09 | Motor region | Right |
| 67.05 | -23.50 | 10.08 | Superior temporal gyrus | Right |
| 63.65 | -33.23 | -6.22 | Middle temporal gyrus | Right |
| 56.77 | -56.73 | 27.37 | Temporoparietal junction | Right |
| -21.47 | 64.43 | 7.44 | Frontal polar | Left |
| -36.69 | 42.16 | 32.13 | Dorsal lateral prefrontal cortex | Left |
| -26.40 | 35.54 | 51.92 | Frontal eye field | Left |
| -51.17 | 38.81 | 6.15 | Inferior frontal gyrus | Left |
| -60.20 | -0.25 | 33.09 | Motor region | Left |
| -67.05 | -23.50 | 10.08 | Superior temporal gyrus | Left |
| -63.65 | -33.23 | -6.22 | Middle temporal gyrus | Left |
| -56.77 | -56.73 | 27.37 | Temporoparietal junction | Left |

2. **The behavior categories in puzzle solving**

We coded every interactive behavior of the dyads during puzzle solving based on Dyadic Parent-Child Interaction Coding System (DPICS) (5,6). Below is a summary of these behavioral categories, excluding physical behaviors, based on their definitions and examples:

| **Table S2.** The DPICS behavior categories. | | | |
| --- | --- | --- | --- |
| Behavior Name(Abbreviation) | Initiator | Definition | Example |
| Negative Talk(NTA) | Parent | The expression of disapproval or rude speech | "I don’t like your attitude." |
| Direct Command(DC) | Parent | A declarative statement containing an explicit order for the child to act | "Put the crayon down." |
| Indirect Command(IC) | Parent | A suggestion phrased as a question to prompt child behavior | "Can you give me a red one?" |
| Labeled Praise(LP) | Parent | Positive evaluation of a specific child attribute or behavior | "You made a fantastic airplane!" |
| Unlabeled Praise(UP) | Parent | General and positive evaluation without specificity | "Great job!" |
| Information Question(IQ) | Parent | A question requiring detailed information | "What do you want to play?" |
| Descriptive Question(DQ) | Parent | A question prompting a yes/no response | "Do you want to play blocks?" |
| Reflection(RF) | Parent | Repetition or paraphrasing of the child’s statement | Child: "It’s blue." Parent: "Blue." |
| Behavior Description(BD) | Parent | Declarative statements describing the child’s observable behavior | "You are drawing a butterfly." |
| Negative Talk(NTA) | Child | The expression of disapproval or rude speech | "I don’t like this game." |
| Command(CM) | Child | Telling or asking the parent to perform an action | "Give me that one!" |
| Question(QU) | Child | Inquiry directed at the parent without suggesting compliance | "Is it special time?" |
| Prosocial Talk(PRO) | Child | Positive statement enhancing interaction | "I like playing special time with you, Mom." |
| Answer(AN) | Child | Appropriate response to a parent’s question | Parent: "What is this?" Child: "A bear!" |
| No Answer(NA) | Child | Failure to respond to a parent’s question | Parent: "What is this?" Child: (no response) |
| No Opportunity to Answer(NOA) | Child | Insufficient time to answer before parent intervenes | Parent: "What is this?" Parent: "It’s a bear!" |

3. **Language characteristics of children**

We analyzed the language characteristics of children during free talking with open source software Computerized Language Analysis (CLAN) (7). For each episode of conversation, we extracted 14 features. Their meanings are listed below:

| **Table S3 The children’s language features during free talking** | |
| --- | --- |
| Feature name | Meaning |
| MLU_Utts | The total number of utterances in the conversation |
| MLU_Words | Mean number of words in each utterance |
| FREQ_types | Number of unique words |
| FREQ_tokens | Total number of words |
| FREQ_TTR | The ratio of the number of unique words and the number of total words |
| %_Noun | The proportion of nouns in total words |
| %_Verb | The proportion of verbs in total words |
| %_pre | The proportion of preposition in total words |
| %_adj | The proportion of adjectives in total words |
| %_adv | The proportion of adverbs in total words |
| %_conj | The proportion of conjunctions in total words |
| %_pro | The proportion of pronouns in total words |
| open_closed | The ratio of the number of open-class words (nouns, verbs, adjectives and adverbs) and the number of closed-class words (prepositions, conjunctions and pronouns) |
